# *FishPi*: a bioinformatic prediction tool to link piRNA and transposable elements

**DOI:** 10.1101/2024.09.10.612046

**Authors:** Alice M. Godden, Benjamin Rix, Simone Immler

## Abstract

**Background:** Piwi-interacting RNAs (piRNA)s are non-coding small RNAs that post-transcriptionally affect gene expression and regulation. Through complementary seed region binding with transposable elements (TEs), piRNAs protect the genome from transposition. A tool to link piRNAs with complementary TE targets will improve our understanding of the role of piRNAs in genome maintenance and gene regulation. Existing tools such as *TEsmall* can process sRNA-seq datasets to produce differentially expressed piRNAs, and *piRScan* developed for nematodes can link piRNAs and TEs but it requires knowledge about the target region of interest and works backwards.

**Results:** We have therefore developed *FishPi* to predict the pairings between piRNA and TEs for available genomes from zebrafish, medaka and tilapia, with full user customisation of parameters including orientation of piRNA, mismatches in the piRNA seed binding to TE and scored output lists of piRNA-TE matches. *FishPi* works with individual piRNAs or a list of piRNA sequences in fasta format. The software focuses on the piRNA-TE seed region and analyses reference TEs for piRNA complementarity. TE type is examined, counted and stored to a dictionary, with genomic loci recorded. Any updates to piRNA-TE binding rules can easily be incorporated by changing the seed-region options in the graphic user-interface. *FishPi* provides a graphic interface using *tkinter* for the user to input piRNA sequences to generate comprehensive reports on piRNA-TE interactions. *FishPi* can easily be adapted to genomes from other species and taxa opening the interpretation of piRNA functionality to a wide community.

**Conclusions:** Users will gain insight into genome mobility and *FishPi* will help further our understanding of the biological role of piRNAs and their interaction with TEs in a similar way that public databases have improved the access to and the understanding of the role of small RNAs.

## Background

Environmental changes and related biotic and abiotic stressors can disrupt the balance and expression of small RNAs and transposable elements (TEs) in the genome ^1^. TEs are mobile DNA elements that can cause insertions in the genome, as well as deletions and genomic rearrangements, initially identified by McClintock in maize but later found to be present across taxa ^2,3^. The mechanisms underlying TE transposition in zebrafish can be divided into two categories: retrotransposons that synthesise an RNA intermediate to insert themselves into the host genome ^4^ and DNA transposons that use a cut-and-paste mechanism to insert themselves into the host genome ^4-6^. Therefore, knowing the family of TE that is mobile can improve our understanding of the mode of TE action and impact on the genome.

Piwi-interacting RNAs (piRNAs) have evolved to repress the action of TEs and constitute a substantial fraction of small RNAs ^7^. The main function of piRNAs is to silence TEs and repetitive elements in animal germ cells to preserve the integrity of the germline genome ^8^. In zebrafish *Danio rerio*, piRNAs are 100x more frequent than miRNAs in zebrafish ovaries and testes ^7^, and small RNAs post-transcriptionally regulate gene expression and regulation ^9^ and the loss of Ziwi leads to apoptosis of germ cells in embryonic development ^7^. Ziwi and Zili proteins provide a piRNA amplification system that is highly conserved. This is important as the availability of complementary piRNAs is essential for survival of germ cells ^7^. In the nematode *Caenorhabditis elegans*, loss of TE silencing by piRNAs can lead to sterility ^10^.

TEs are not randomly distributed in the genome but they have preferred regions for insertion, including distorted and bent DNA ^5,11^, or open chromatin regions ^11,12^. Vast populations of piRNAs are complementary to TE messenger RNAs (mRNA) ^13^. In *C. elegans*, the 2^nd^ to the 7^th^ nucleotide at the 5’ end of the piRNA ^14^ or the 2^nd^ to the 8^th^ nucleotide ^15^ have both been suggested to be the piRNA-TE seed region. Similarly, in the house mouse *Mus musculus*, the 2^nd^ to 8^th^ nucleotides have been identified as a seed region ^16^. In contrast to mammals, most zebrafish piRNAs map to TEs, and the LTR families in particular ^7^. In zebrafish and other teleost species, the first ten nucleotides at the 5’ end of the piRNA are loaded into the Argonaute machinery to produce Ziwi (zebrafish Piwi)-bound piRNAs complementary to TE transcripts ^7,13^. The same is true for many other organisms including *Drosophila*, as piRNAs show complementarity to other antisense piRNAs in their first ten nucleotides, which associate with PIWI proteins to generate the mature 5’ end of the piRNA ^7^ forming part of the ping-pong amplification cycle where piRNAs act and bind complementarily to transposons ^17^. It is the PIWI-bound piRNA with the piRNA seed that recognises a complementary region on the transposon and leads to silencing ^18,8^. In *Drosophila*, piRNAs cleave their target mRNAs (TEs) after the 10^th^ nucleotide from 5’ end of the piRNA^19^. Given the evolutionary conservation of piRNA biogenesis and TE silencing mechanisms ^7,20^, using the first ten nucleotides of piRNA as a putative piRNA seed region is what we use as a starting point. However, *FishPi* allows users to propose custom piRNA seed regions to predict piRNA:TE activity in the event of further work confirming or changing this prediction.

TEs are known to be activated through specific environmental stressors. In *C. elegans* mutant strains lacking the ability to generate piRNAs, heat-treated worms displayed transgenerational impairments for up to three subsequent generations with thermal stress reducing the rate of piRNA biogenesis ^21,22^. In *Drosophila*, the *mariner-Mos1* transposon was enriched following heat-stress but not following ultraviolet irradiation ^23^. TEs are expressed in a tissue-specific manner and can affect transcription and gene expression ^24^ by generating new genes and mRNAs through genomic rearrangements and *de novo* mutations, and cause alteration of regulatory networks and even trigger immune responses ^5^. Understanding the interaction between piRNAs and their targets is therefore key to understand the defence mechanisms in the germ line against the effects of environmental conditions.

We developed the software *FishPi* to enable mapping zebrafish piRNAs against complementary TEs and link them to understand their biological association ^13^. *FishPi* facilitates research into the function of piRNAs and accelerates analyses in the same way as microRNA databases like *Targetscan* have supported small RNA research ^25^. *FishPi* provides a user-friendly graphic interface (Figure 1) and generates a comprehensive output that can be exported as a report including complementary TE counts, family level annotation, chromosomal and counts plots, with the option to export the full list of complementary TEs and their sequences. The user can enter individual piRNAs or lists of piRNAs in fasta format. For zebrafish and other model organisms, a list of piRNA sequences are available at piRBase (http://bigdata.ibp.ac.cn/piRBase/browse.php), ^26^ or piRNAclusterDB (https://www.smallrnagroup.uni-mainz.de/piRNAclusterDB/) ^27^. *FishPi* can easily be used for any species for which the relevant information such as a high-resolution reference genome and sufficient piRNA sequences are available.

**Figure 1.**
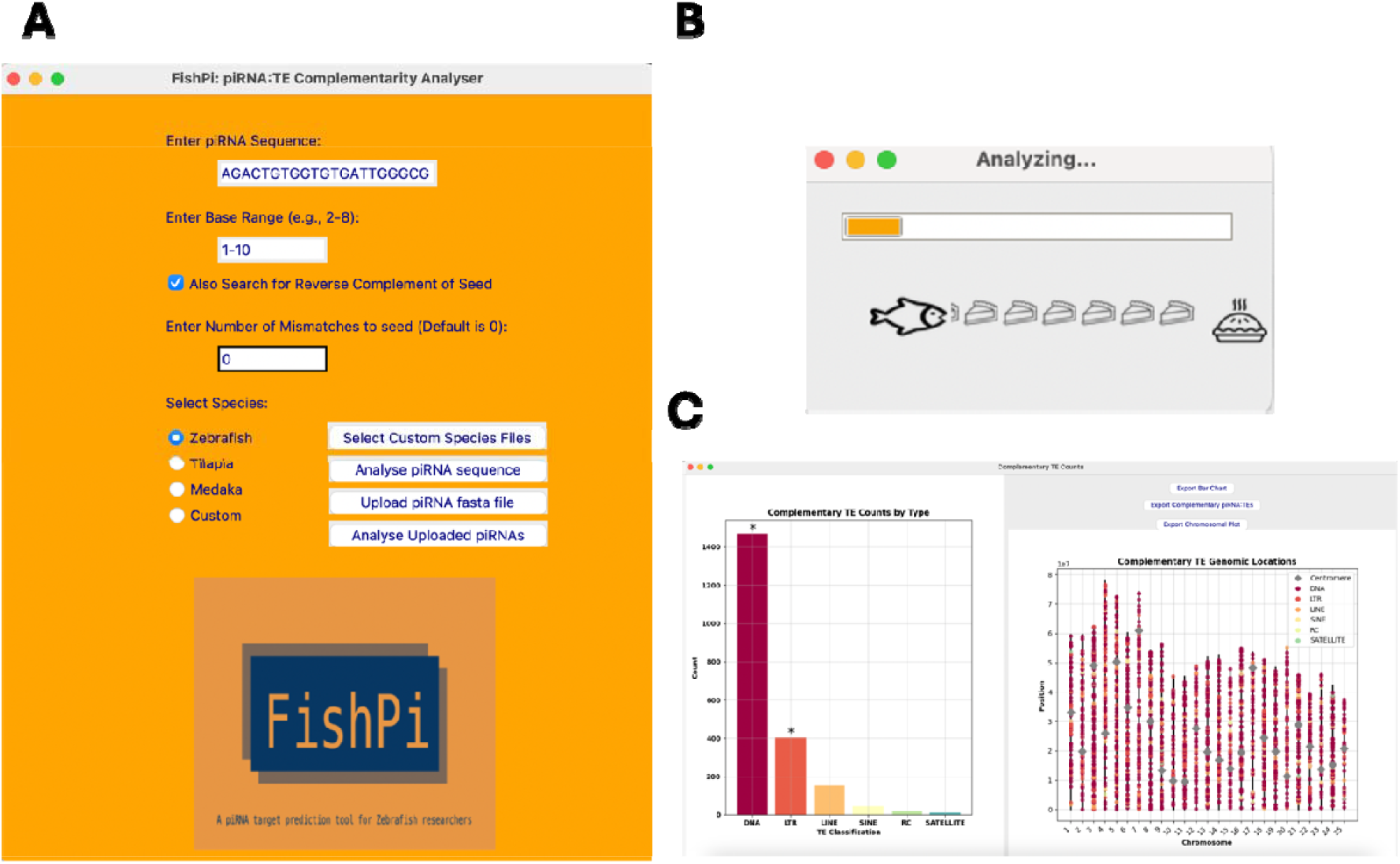
The graphic user interface of FishPi with and pop-out window with exportable results for dre-piRNA 5’-TACACGAAGACTGTGGTGTGATTGGGCG-3’. The buttons included allow users to export the plots at hi-resolution for presentation and publication. Additionally, the user may export a .csv file list of all complementary TE mRNAs for further downstream analysis.

### Implementation

*FishPi* is a Python-based programme and uses the following packages and versions: Python v3.11, Tkinter (included in Python v3.11 ^28^), Pillow 10.0.1 ^29^, and Matplotlib 3.8.0 ^30^. The code is platform independent (see Figure 2 for overview of processes) and follows the installation guidance for Conda on Github (https://github.com/alicegodden/fishpi ). The TE families assigned to each class were checked using RepeatMasker and Dfam (Table 1) ^31^. Input TE fasta files can be found on Zenodo: https://zenodo.org/records/13975588.

**Table 1.** Transposable element families assigned to transposable element class.

| **Class** | **Transposable element family** |
| --- | --- |
| DNA | hAT, Tc1, Tc-Mar, Harbinger, Enspm, Kolobok, Merlin, Crypton, PiggyBac, Dada, Zatar, Ginger, TDR, Polinton, Maverick, Acrobat, Looper, TZF, Angel, Mariner |
| LTR | Gypsy, DIRS, Ngaro, ERV, Pao, Copia, BEL, HERV, Bhikari |
| LINE | L1, L2, L1-Tx1, Rex-Babar, RTE, Penelope, Keno, Rex |
| SINE | Alu, tRNA-V-RTE |
| RC | Helitron |
| Satellite | BRSATI, MOSAT |

**Figure 2.**
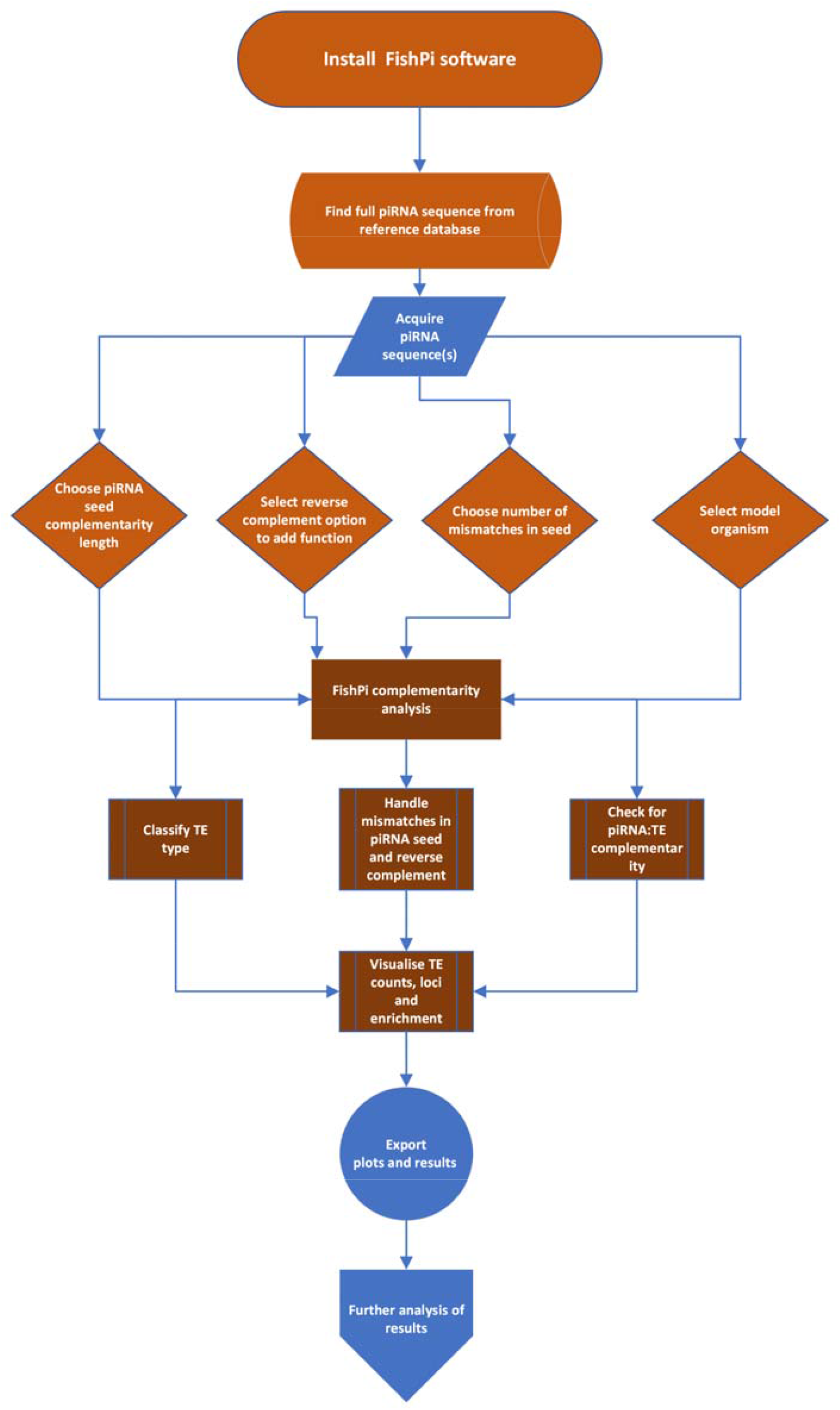
Flow chart covering the key steps and processes of FishPi.

#### Input file preparation

To generate a fasta file of the TE sequences from the zebrafish reference genome GRCz11, a TE.bed file with clade was created by using the UCSC table genome browser ^32^ as follows: Vertebrate, group: Variation and Repeats, genome: zebrafish, assembly: May 2017 GRCz11, track: RepeatMasker, table: rmsk. Reference genome files for zebrafish, medaka and tilapia were obtained from Ensembl. Genomes were decompressed with *gunzip* and indexed with *samtools faidx refgenome*.*fa*. A gtf TE annotation file was generated with: *perl makeTEgtf*.*pl -c 6 -s 7 -e 8 -o 10 -t 11 fish_rmsk > fish_TE*.*gtf*. This script was obtained from the Hammel lab (https://labshare.cshl.edu/shares/mhammelllab/www-data/TEtranscripts/TE_GTF/makeTEgtf.pl.gz ) ^33^. Only necessary columns were retained with following command in fish_TE.gtf: *cut -f1*,*2*,*3*,*4*,*5*,*7*,*8*,*9*,*10*,*11*,*12*,*13*,*14*,*15*,*16 fish_TE*.*gtf > fish_TE*.*use*.*gtf*. The awk script to convert .gtf to .bed file can be found on the README.md https://github.com/alicegodden/fishpi/blob/main/README.md , along with *sed* commands to handle different chromosome naming conventions. Finally, the steps to extract DNA sequences from the reference genome are: *bedtools getfasta -s -name -fi Danio_rerio*.*GRCz11*.*dna*.*primary_assembly*.*fa -fo GRCz11*.*teseqs*.*use*.*fasta -bed GRCz11*.*teannotation*.*bed*.

To generate chromosome length files for chromosomal plots “species_chrom_end.txt” files, the user needs to navigate to the UCSC table browser (https://genome.ucsc.edu/cgi-bin/hgTables) and select the species and reference genomes as above. Under the “Group” drop down menu, select “*All Tables*”. Under the “*Table*” drop down menu select “cytoBandIdeo”. To download the file in .txt format, the first column needs to be chromosome names/numbers and second column to be the length of each chromosome. The function that represents the inclusion of custom species files is *select_custom_files()*.

Users can choose their own piRNA sequence(s) through individual pasting of piRNA sequences or upload a list of named piRNAs, *upload_pirna_file*. The piRNA seed region sequence can be specified. For teleosts, we recommend 1-10 nucleotides (default setting). “*Reverse complement*” can be checked to add the reverse of piRNA seed for complementarity matching. Mismatches in the piRNA seed can be included, the number of these can be specified, where zero is the default setting. Once executed with *analyse_sequence()* or *analyse_uploaded_pirna()*, complementary TEs are identified in any orientation with *is_complementary()* and with mismatches with *introduce_mismatches()*. Visualisations are provided by the function *create_results()*. The resulting bar chart includes a binomial distribution test, with stars on the bar chart denoting *P* _≤_ 0.05. This is to show that the likelihood of a specific TE family is complementary to the piRNA at a significantly higher than expected ratio. The functions *classify_te_type()* group complementary TEs by family for plotting and *plot_te_chromosomal_locations()* displays a chromosomal plot of complementary TE loci in the genome. All results can be exported with *export_complementary_te_list()*, a .csv output file of results including piRNA names (when fasta file of piRNAs is uploaded), TE name and location, orientation of complementarity and complementarity percentage of piRNA (not including seed region) to TE. The functions *export_bar_chart()* and *export_chrom_plot()* allow users to save high resolution images for presentation and publication.

## Results & Discussion

*FishPi* is designed for users with minimal coding experience. A detailed manual can be found in the README on the git repository: https://github.com/alicegodden/fishpi/blob/main/README.md. The simple graphic user interface offers a point and click tool to rapidly gain insight into many piRNA sequences. Each time a new piRNA sequence is provided, the programme resets itself before providing the new results. As more research into piRNA-TE seed sites evolve, users can change their search criteria. Additionally, users can generate and use other reference TE genome files and use *FishPi* for other model systems through the custom options. This makes *FishPi* particularly powerful for users looking at non-model systems. The software has a graphic user interface for *FishPi* (Figure 1A), and the outputs can be exported as high-resolution figures presenting raw count data of the different target TE categories for presentation (Figure 1B). Additionally, users can export a .csv file with all complementary TE mRNA sequences, along with genomic co-ordinates, allowing the user to conduct further downstream analyses and explore each class of TE individually. *FishPi*’s advantage is that as more research into piRNA-TE seeds are validated, the user can customise their search and input different seed regions for analysis. Additionally, users can use this for any species and there is option for custom species input.

Tools for the study of piRNAs and tools for piRNA target analyses do exist but have limited application bandwidths. *PiRNAQuest* V.2 for example is an online tool to annotate and identify piRNA clusters and it can identify piRNA targets, but these are limited to mouse and diseases related to cancers ^34^. *piRPred* is piRNA identification tool based on an identification algorithm with k-nearest neighbours’ kernel to improve piRNA classification ^35^. The tool *miRanda* is primarily designed for miRNA gene target prediction to look at piRNA target genes, rather than TEs ^36,37^. *FishPi* fills a current gap and offers a user-friendly, adaptable and customisable interface for identifying transposons targeted by piRNAs.

## Conclusions

Currently available tools to link piRNA and TEs are limited and require knowledge of known targets or are restricted to a pre-defined genome and species due to piRNA binding rules ^14-16,38^. *FishPi* is ideal for users with a list of candidate piRNAs. This is tool that can easily be adapted to any organism in a user-friendly GUI format. *FishPi* is fully adaptable and customisable so will continue to be user-friendly and useable as our understanding of piRNA targeting improves. It generates detailed reports and files that can be used for further analysis including statistical analysis of TE family targets in enrichment bar charts, loci analysis in chromosomal plots, and exportable spreadsheets with piRNA:TE targets with scoring via percentage complementarity for further filtering. *FishPi* software is freely available and is distributed under GPL-3.0 licence.

## Availability and requirements

**Project name:** FishPi

**Project home page:** https://github.com/alicegodden/fishpi

**Operating system(s):** Platform independent

**Programming language:** Python

**Other requirements:** Python v3.11

**License:** GPL-3.0

**Any restrictions to use by non-academics:** GPL-3.0 license needed

## Declarations

### Ethics

No ethics approvals were required for this research project.

### Consent for publication

All authors consent to publication.

### Availability of data and materials

*FishPi* is available on the Git repository here: https://github.com/alicegodden/fishpi

### Competing interests

Authors declare no financial or competing interests.

### Funding

This project was funded by a Consolidator Grant from the European Research Council to SI (SELECTHAPLOID – 101).

### Author’s contributions

AMG conceived the idea for the project, AMG and BR wrote the software, BR tested the software, AMG wrote the manuscript and SI contributed to the writing and provided guidance throughout.

## Acknowledgements

AMG would like to thank Immler lab colleagues for testing FishPi.

## List of abbreviations

mRNA: Messenger RNA
PiRNA: Piwi-Interacting RNA
TE: Transposable element

## Notes

https://github.com/alicegodden/fishpi

### Competing Interest Statement

The authors have declared no competing interest.

### Summary of Updates

This version of the manuscript has been revised to update the improved functionality and as such user instructions for installation and use of FishPi.

